# Langerhans cells regulate immunity in adulthood by regulating postnatal dermal lymphatic development

**DOI:** 10.1101/2024.07.12.603312

**Authors:** Ji Hyun Sim, Richard Bell, Zhonghui Feng, Susan Chyou, William D Shipman, Raghu P. Kataru, Lionel Ivashkiv, Babak Mehrara, Theresa T. Lu

## Abstract

The communication between skin and draining lymph nodes is crucial for well-regulated immune responses to skin insults. The skin sends antigen and other signals via lymphatic vessels to regulate lymph node activity, and regulating dermal lymphatic function is another means to control immunity. Here, we show that Langerhans cells (LCs), epidermis-derived antigen-presenting cells, mediate dermal lymphatic expansion and phenotype acquisition postnatally, a function is independent of LC entry into lymphatic vessels. This postnatal LC-lymphatic axis serves in part to control inflammatory systemic T cell responses in adulthood. Our data provide a tissue-based mechanism by which LCs regulate T cells remotely across time and space and raise the possibility that immune diseases in adulthood could reflect compromise of the LC-lymphatic axis in childhood.

## INTRODUCTION

The communication between skin and draining lymph nodes is crucial for timely and well-regulated immune responses to skin insults. Skin-derived antigens, cells, and signals that are brought by dermal lymphatic vessels to draining lymph nodes act directly on lymph node cells to regulate immunity. Regulating how well lymphatic vessels function provides an additional means of communication that is indirect and that allows local skin events to remotely control downstream lymph node activity. The impact of regulating lymphatic function on immune responses can be seen in mouse models with a dearth of dermal lymphatic vessels. These models show reduced responses to soluble antigens, reduced tolerogenic mechanisms, and spontaneous autoimmunity (*1–5*). In humans, evidence of lymphatic flow dysfunction is found in autoimmune diseases (*6, 7*), and correction of lymphatic flow in disease models leads to reduced lymph node B cell responses (*7*). Insights into local skin mechanisms that can regulate dermal lymphatic function, then, provides opportunities for better understanding and treatment of immune diseases.

Langerhans cells have unique properties among myeloid cells, long considered to be a prototypic example of a DC migrating from skin to draining lymph nodes to present antigen to T cells but, in more recent years, found ontogenetically to be derived from embryonic precursors of tissue-resident macrophages (*8–11*). Residing and self-renewing in the epithelium and migrating constitutively into the dermis and, via dermal lymphatics, to draining lymph nodes (*11–13*), LCs can have regulatory roles in immunity, with Langerin-DTA (Lang-DTA) mice that constitutively lack LCs showing enhanced contact hypersensitivity (CHS) responses (*14*). This regulatory effect was attributed to increased T cell priming in draining lymph nodes during the sensitization phase and dependent on LC MHCII expression, and increased CHS responses could be replicated by acute LC ablation (*14, 15*) (*16*), all suggesting that LC regulatory functions were related to their roles as antigen-presenting cells. However, Lang-DTA mice also showed greater skin inflammation to locally injected Candida albicans, a process that was independent of adaptive immunity (*17*). This latter result suggested that at least some of the regulatory effects of LCs may be local to the skin and independent of antigen-presenting function, but mechanisms by which LCs are immune regulatory at the level of the skin and how that translates into lymph node regulation remains poorly understood.

We and others have more recently shown that LCs have tissue-modulating functions. In studying mechanisms of lupus photosensitivity whereby patients can develop inflammatory skin lesions with even ambient ultraviolet radiation (UVR), we showed that LCs protect epidermal integrity by limiting UVR-induced keratinocyte death (*18*). LCs express ADAM17, a metalloprotease that clips and releases EGFR ligands from the membrane, allowing the EGFR ligands to bind keratinocyte EGFR and promote cell survival. Multiple SLE models have shown defective LC ADAM17 activity and consequently greater UVR-induced skin inflammation, which can be ameliorated by circumventing the LC ADAM17 dysfunction by topical EGFR ligand (*18, 19*). Horsley and colleagues have also shown that LCs express high levels of VEGF- A and other angiogenic factors and demonstrated a role for LCs in blood vessel angiogenesis during wound healing (*20*). During their journey from the epidermis to draining lymph nodes, LCs interact directly with lymphatic vessels as they migrate toward and enter the vessels and are thus well positioned to potentially regulate lymphatic function. The tissue-modulating functions of LCs and their interactions with lymphatic vessels led us to examine their role in regulating dermal lymphatics and the implications for controlling immunity.

## RESULTS

### Constitutive lack of LCs alters lymphatic endothelial cell (LEC) numbers, LEC phenotype, and lymphatic function in adult skin

We examined skin lymphatic endothelial cell (LEC), blood endothelial cell (BEC), and CD45- non-endothelial (EC) stromal cell numbers in 8 week old wild-type (WT) and Lang-DTA mice. Consistent with previous characterization(*14, 21*), Lang-DTA mice lacked ear skin LCs as early as 7 days of age (**fig. S1A-B**). LCs were also lacking in the auricular lymph nodes that drain the ears in Lang-DTA mice **(fig. S1B)**. Remarkably, Lang-DTA mice showed reduced numbers of LECs in ear skin when compared to WT mice **(Fig. 1A upper row and, for gating scheme, fig. S1C)**. BEC numbers were also reduced in Lang-DTA skin **(fig. S1D, D)**, although the reduction was modest compared to LECs. The more numerous PDPN+ non-EC stromal cells and total skin cellularity were unaffected **(fig. S1C, E-F)**. In contrast to skin, lymph node LECs, BECs, PDPN+ fibroblastic reticular cells (FRCs), and total lymph node cellularity were unaffected **(Fig. 1A bottom row and fig. S1G-J)**. These results showed that a constitutive lack of LCs resulted in reduced dermal LEC numbers.

**Fig. 1.**
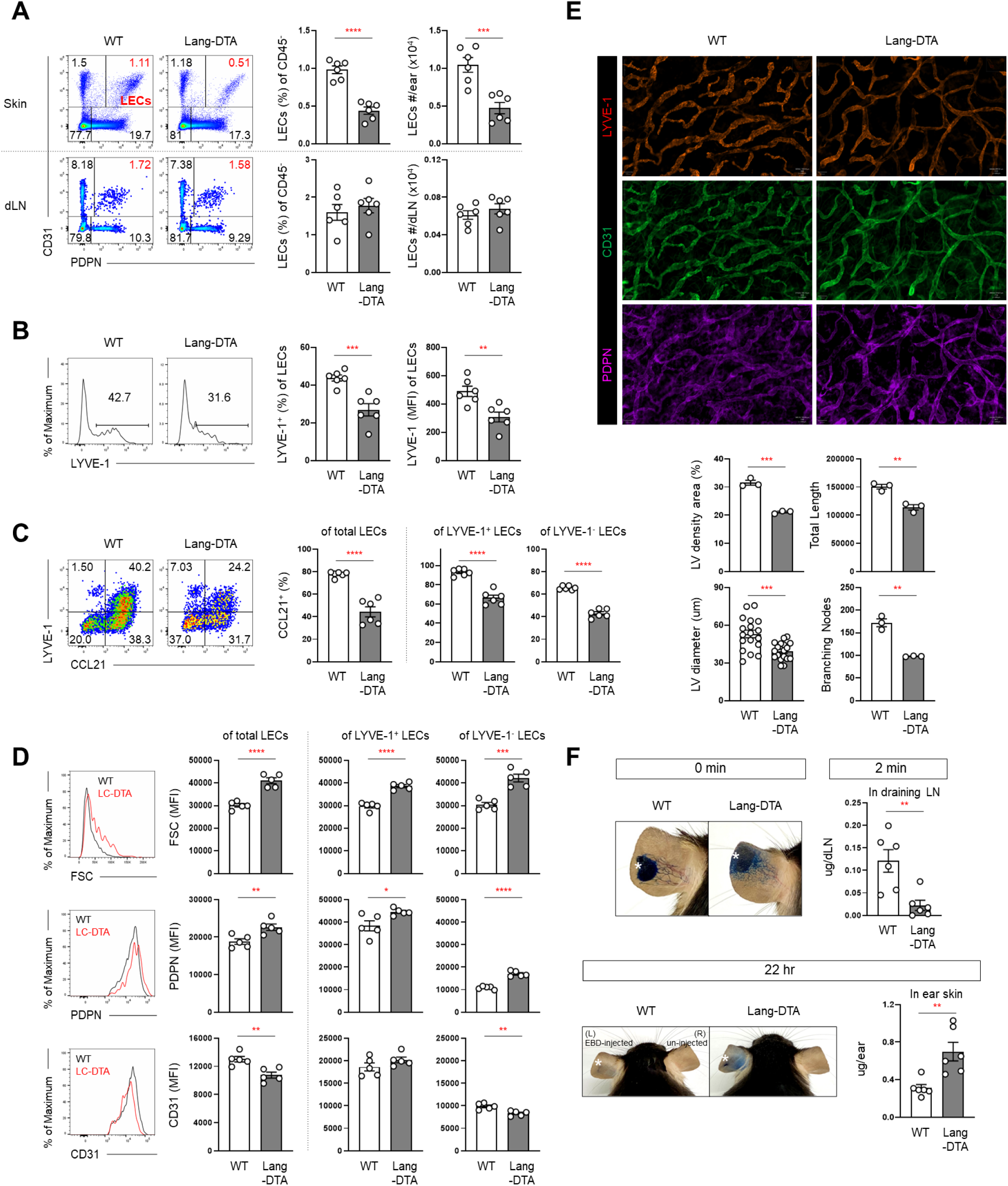
Constitutive absence of LCs alters lymphatic endothelial cell (LEC) numbers, LEC phenotype, and lymphatic vessel function in skin. (**A-D**), Ear skin of 8 week old WT and Lang-DTA mice was enzymatically digested and stained for flow cytometry analysis. (**A**) LEC quantitation in ear skin (top row) and enzymatically digested draining auricular lymph nodes (dLN, bottom row). Representative flow cytometry plots of CD45-cells (left), percentage of CD45-cells and number of LECs (right). (**B**) LEC LYVE-1 expression. Representative histograms (left), percentage that are LYVE-1+ and LYVE-1 mean fluorescence intensity (MFI) (right). (**C**) LEC CCL21 expression. Representative plots (left) and percentage that are CCL21+ of total, LYVE-1^+^, and LYVE-1^-^LECs (right). (**D**) LEC forward scatter (FSC) (top), PDPN (middle), and CD31 (bottom). Representative histograms (left) and MFI of total, LYVE-1^+^, and LYVE-1^-^LECs (right). (**E**) Ears from 8 week old mice were subject to whole mount immunofluorescence staining for LYVE-1 (red), CD31 (green), and PDPN (magenta) and analyzed by whole side imaging. Representative images (top), quantification of lymphatic vessel (LV) density and total length, diameter of each vessel segment, and branching node numbers (bottom). (**F**) Lymphatic drainage assessment. Evans Blue Dye (EBD) was injected at tip of the ear pinna (stars) and indicated tissues were assayed for EBD content at indicated time points. Each symbol represents one mouse (**A-F**) except for (**E**, LV diameter), where each symbol represents one vessel segment. Bars represent the mean and error bars represent s.e.m. *p<0.05, **p<0.01, ***p<0.001, ****p<0.0001 using two-tailed unpaired Student’s t-test. Mice per condition: n=3-6 (**A-F**) and data are from 2 (**D**), 3 (**B- C, E**) and 4 (**A, F**) independent experiments.

Lang-DTA dermal LECs also showed phenotypic alterations, expressing lower levels of the dermal lymphatic capillary marker lymphatic vessel endothelial hyaluronan receptor-1 (LYVE1) (*22*) **(Fig.1B)**. Both LYVE1+ and LYVE1-subsets in Lang-DTA mice expressed less CCL21, a CCR7 ligand that mediates DC migration from the dermis into lymphatic capillaries and then toward the lymph node (*23–25*) **(Fig. 1C)**. Lang-DTA LECs were also larger, as indicated by increased forward scatter (FSC) **(Fig. 1D, top row)**, and expressed increased PDPN **(Fig. 1D, middle row)**, which is upregulated upon cell stress in mesenchymal cells (*26–28*). LYVE1- cells also expressed reduced CD31 **(Fig. 1D, bottom row)**, potentially indicative of greater susceptibility to cell death (*29*). In contrast, skin BEC size and CD31 levels were unaffected **(fig. S1K)**.

Whole mount staining of the skin corroborated the flow cytometry results, showing reduced lymphatic vessel density and total length **(Fig. 1E)**. Vessel diameters were also smaller, and there were fewer branch points, suggesting less complexity of the vascular network **(Fig. 1E)**. These results together indicated that Lang-DTA skin had fewer LECs with altered phenotype and fewer and smaller lymphatic vessels.

We performed Evans blue dye (EBD) lymphangiography to assess for lymphatic vessel integrity and flow. In contrast to the sharply delineated lymphatic vessels in WT mice, Langerin- DTA ears showed a diffuse distribution of EBD immediately after injection **(Fig. 1F, 0 min and fig. S2A, left panels)**, suggesting leakage of EBD from the vessels and reduced flow from the skin. Consistent with the skin findings, less EBD accumulated in draining lymph nodes at 2 minutes after EBD injection and there was less clearance of EBD from the ears at 22 hours **(Fig. 1F, 2 min, 22 hr, and fig. S2B, left panels)**. The compromised lymphatic drainage was not unique to ear skin, as EBD accumulation in popliteal lymph nodes after footpad inject was also reduced in Lang-DTA mice **(fig. S2C).** These results indicated reduced lymphatic drainage function in the skin of Langerin-DTA mice. Together, the results suggested that the constitutive absence of LCs compromised LEC and lymphatic vessel numbers and phenotype and lymphatic flow function in the skin.

### LCs are required during postnatal dermal lymphatic expansion

We asked the extent to which the dermal LEC abnormalities in adult Langerin-DTA mice reflected developmental issues. While WT LECs steadily expanded in numbers between 2 and 8 weeks of age, Lang-DTA LECs showed an early expansion but then failed to expand after 4 weeks **(Fig. 2A)**. This resulted in a small but detectable decrease in LEC numbers when compared to WT mice by 5 weeks of age that became more dramatic over time **(Fig. 2A)**. Similarly, Lang-DTA LEC LYVE1, CCL21, PDPN, and CD31 expression were unchanged from WT levels at 3 weeks but showed alterations by 4-6 weeks of age **(Fig. 2B-C and fig. S2D)**. EBD assays and whole mount staining confirmed normal lymphatic vessel density and flow function at 4 weeks of age **(fig. 2A-C,E-F)**. These data indicated that the compromised lymphatic expansion and phenotypes seen in adult Lang-DTA mice reflected compromised postnatal lymphatic development.

**Fig. 2.**
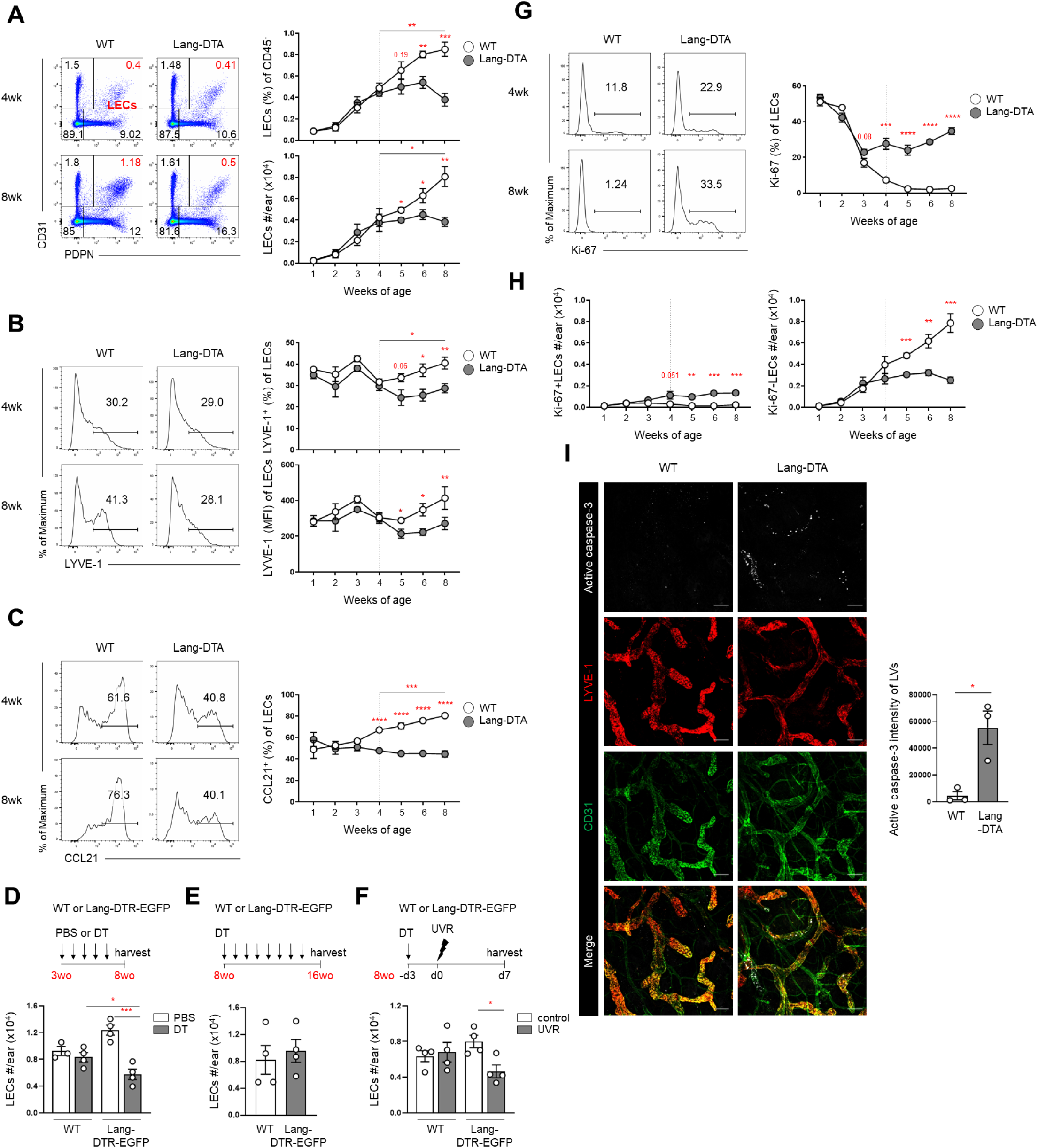
LCs are required for postnatal lymphatic development and, in adulthood, lymphatic maintenance during inflammation. (**A-C**) Ear skin from WT and Lang-DTA at indicated ages was examined by flow cytometry. (**A**) LEC quantitation. Representative plots of CD45-cells (left), percentage and number of LECs (right). (**B**) LEC LYVE-1 expression. Representative histograms (left), percentage and LYVE-1 MFI (right). (**C**) LEC CCL21 expression. Representative histogram (left) and percentage (right). (**D-F**) WT and Lang-DTR-EGFP mice at indicated weeks of age (weeks old (wo)) were treated with PBS or diphtheria toxin (DT) weekly (**D, E**) or once (**F**) and ear LEC numbers evaluated by flow cytometry. For (**F**), mice were exposed to ultraviolet radiation (UVR) at day 0. Experimental setup (top), LEC numbers (bottom). (**G, H**) Ear skin from WT and Lang-DTA mice at indicated ages was examined for LEC proliferation, as indicated by ki67 staining. g, Representative histogram of LEC Ki-67 (left) and percentage (right). h, Ki-67+ (left) and Ki-67-(right) LEC numbers. (**I**) Ears from 8 week old mice were subject to whole mount immunofluorescence staining for activated caspase-3 (white), LYVE-1 (red), and CD31 (green) and analyzed by confocal microscopy. Representative images (left), intensity of active caspase-3 in CD31+LYVE-1+ vessels (right). For (**A-C, G-H**), each symbol represents average of 4-7 mice per condition at each time point; for (**D-F, I**), each symbol represents one mouse. Bars represent the mean and error bars represent s.e.m. For (**A-C, G-H**), WT and Lang-DTA values were compared statistically at each time point. *p<0.05, **p<0.01, ***p<0.001, ****p<0.0001 using two-tailed unpaired Student’s t-test. Mice per condition: n=4-7 (**A-C, G-H**) , 3-4 (**D**), 4 (**E-F**), 3 (**I**) and data are from 2 (**E, I**), 3 (**A-C, F-H**), and 4 (**D**) independent experiments.

We sought to better understand when LCs were needed for lymphatic expansion and to ask about their role in lymphatic maintenance in adulthood. Depleting LCs from 3 weeks of age in Lang-DTR-EGFP mice that allow for inducible Langerin+ cell (LCs and Langerin+ cDC1) depletion upon DT treatment (*21, 30–32*) led to reduced LEC numbers by 8 weeks of age **(Fig. 2D)**. In contrast, DT treatment of Lang-DTR-EGFP mice from 8 weeks of age onward for 2 months had no effect on LEC numbers **(Fig. 2E)**. These results using a 2^nd^ model to deplete LCs further supported a role for LCs in lymphatic expansion and suggested that LCs are required postnatally between 3 and 8 weeks of age for LEC expansion but not during adulthood for LEC maintenance.

Because inflammatory processes can recapitulate ontogenic processes, we asked about the role of LCs in adulthood in response to an inflammatory injury. Ultraviolet radiation exposure did not affect control mice but led to a reduction in LEC numbers in DT-treated Lang-DTR-EGFP mice **(Fig. 2F)**, suggesting that, in adulthood, LCs are required to maintain LECs during inflammation.

We examined for altered LEC proliferation or apoptosis rates that could result in the impaired postnatal expansion in Langerin-DTA mice. While WT LECs showed a steady decline in proliferation rate from 1 to 5 weeks of age, Lang-DTA LECs showed an abrupt cessation of the decline at 4 weeks of age, resulting in a dramatically elevated proliferation rate from 4 weeks onward **(Fig. 2G).** Absolute numbers of ki67+ LEC numbers were higher in Lang-DTA mice while there was a decline in ki67-cells **(Fig. 2H)**, suggesting that the failure of LECs to expand in number reflected a loss of non-proliferating cells. Indeed, 8 week old Lang-DTA mice showed elevated activated caspase 3 in vessels when compared to WT mice **(Fig. 2I),** suggesting greater apoptosis. Together, these results suggested that the lack of LEC expansion in Lang-DTA mice reflected compromised survival of non-proliferating cells.

In contrast to dermal LECs, dermal BECs, and PDPN+ non-EC stromal cells did not show as robust expansion between 4 and 8 weeks of age **(fig. S3A)**. Dermal BECs did show reduction of FSC, CD31, and proliferation rates, similar to although with a lesser magnitude than LECs **(fig. S3B)**. In lymph nodes, LECs were different from skin LECs in showing no expansion in numbers, and downregulating rather than upregulating LYVE1 **(fig. S3C-D)**. Lymph node LECs were similar to skin LECs in upregulating CCL21 **(fig. S3E)** and showing minimal changes in FSC, PDPN, and CD31 levels **(fig. S3F)**. Lang-DTA mice did not differ from WT mice in the pattern of these postnatal changes in skin non-LEC stromal cells and lymph node LECs **(fig. S3A-F)**. Thus, skin LECs showed developmental features distinct from that of other skin stromal cells and lymph node LECs, consistent with the differential dependence on LCs.

### LCs promote LEC expansion directly via VEGF-C and PlGF

We considered the molecular mediators by which LCs regulated LEC expansion. LCs have been shown to express high levels of angiogenic factors VEGF-A and placental growth factor (PlGF) (*20*) (Immgen.org). VEGF-C is a key lymphangiogenic factor for prenatal lymphatic development and, in adulthood, inflammation-associated lymphangiogenesis (*33–37*). (Blood vessel) angiogenic mediator VEGF-A signals through VEGFR1 and VEGFR2, PlGF through mainly VEGFR1, and VEGF-C mainly through VEGFR3 (*35–38*). Treating WT mice from 3 weeks of age for 2-3 weeks with blocking antibodies to VEGFR1, VEGFR2, VEGFR3 singly and in combination showed that blockade of all three VEGF receptors reduced postnatal lymphatic expansion **(Fig. 3A and fig. S4A-C).** Given the high expression of PlGF by LCs, we assessed the combination of PlGF and VEGFR3 blockade, and this was sufficient to partially block postnatal lymphatic expansion (**Fig. 3A and fig. S4C)**, suggesting that PlGF and VEGF-C at least in part mediated the postnatal lymphatic expansion. PlGF and VEGF-C combined were sufficient to rescue postnatal LEC expansion in Lang-DTA mice **(Fig. 3C)**. In vitro, LCs were sufficient to promote LEC survival and did so in an PlGF- and VEGFR3-dependent manner **(Fig. 3C-D)**. Our results suggested that LCs mediated postnatal LEC expansion by a combination of PlGF+VEGF-C and act directly on LECs.

**Fig. 3.**
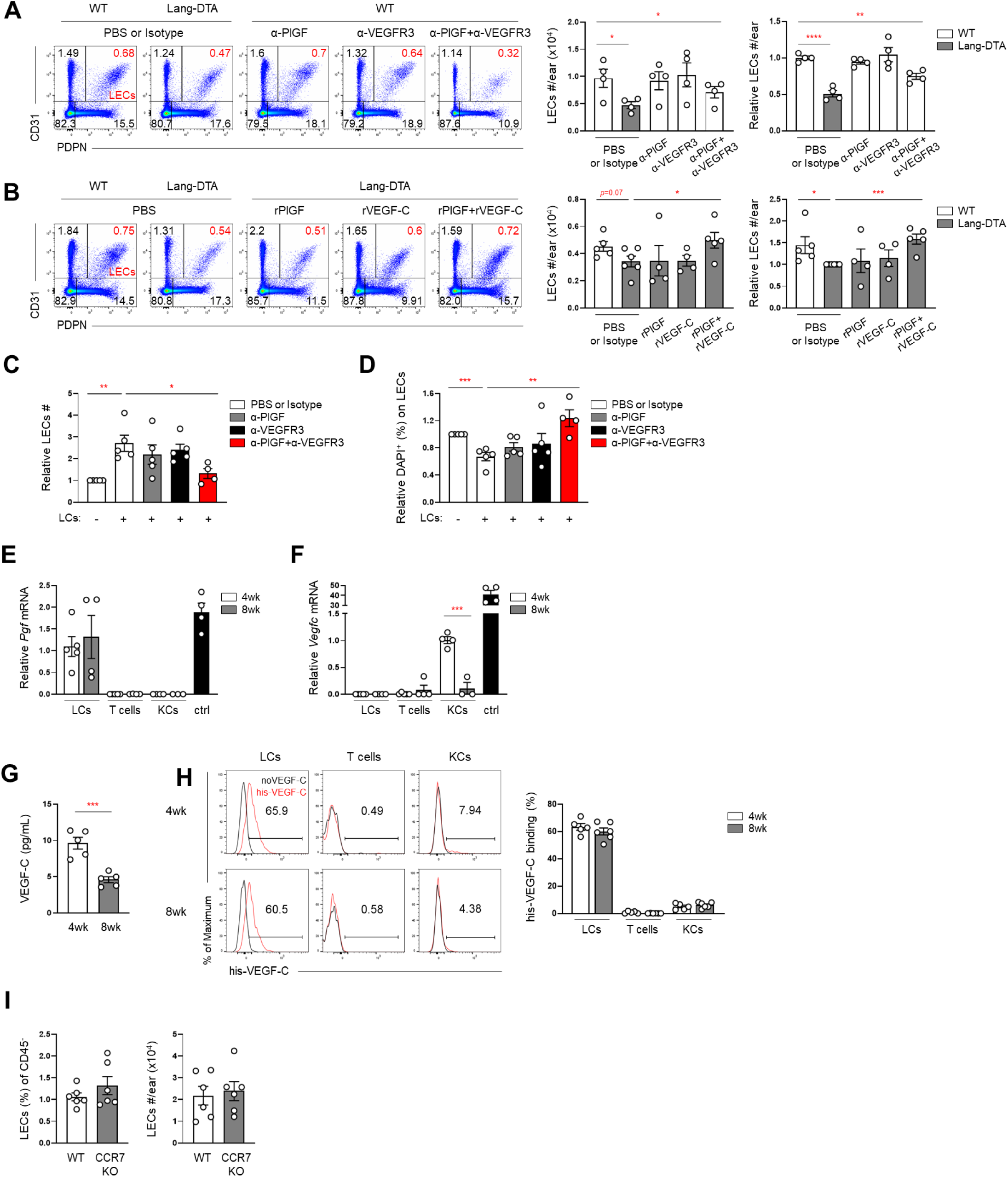
LCs promote LEC development via VEGF-C and PlGF and in a CCR7-independent manner. (**A-B**) 3 week old mice of indicated genotypes were treated intradermally in ear skin with indicated blocking antibodies for 2-3 (**A**) and with indicated cytokines for 2 (**B**) weeks before ear LECs were quantitated. Representative plots (left), absolute and relative LEC numbers (right). (**C, D**) Murine dermal LECs were cultured with or without LCs and with anti-PIGF, anti-VEGFR3, or anti-PIGF+anti-VEGFR3 before LECs were quantitated. Relative LEC numbers (**C**) and percentage that are DAPI+ (**D**). (**E-F**) mRNA expression of Pgf (**E**) and Vegfc (**F**) by sorted LCs, T cells, and keratinocytes (KCs) from 4 and 8 week old WT mice. Positive control (ctrl) was murine placenta. (**G**) LCs from 4 and 8 week old WT mice were cultured for 2 days and supernatant was assessed for VEGF-C by ELISA. (**H**) VEGF-C cell binding. LCs and other cells in the medium after a 48-hour crawl-out from 4 and 8 week old mouse epidermal sheets were incubated with his-tagged VEGF-C which was detected with anti-his tag. Representative histograms (left) and percentage of indicated cells that bound his-VEGF-C (right). (**I**) Ear skin LECs in 8 week old WT and CCR7 KO mice were quantitated. Percentage (left) and number of LECs (right). Each symbol represents one mouse. Bars represent mean and error bars s.e.m. *p<0.05, **p<0.01, ***p<0.001, ****p<0.0001 using two-tailed paired (**A**, left) or unpaired Student’s t-test (**A**, right, **B-I**). Mice per condition: n= 4 (**A**), 4-6 (**B**), 4-5 (**C-D**), 3-5 (**E-F**), 5 (**G**), 5-6 (**H**), 6 (**I**) and data are from 2 of multiple similar experiments (**H**) and 3 (**A,C-F**), 4 (**G, I**) and 5 (**B**) independent experiments.

We assessed for LC expression of PlGF and VEGF-C at different ages. Quantification of LC *Pgf* mRNA showed expression at levels comparable to that found in the placental without changes with age **(Fig. 3E)**. *Vegfc* mRNA could not be detected in LCs, but keratinocytes were found to express *Vegfc,* and these levels dropped dramatically between 4 weeks of age and adulthood **(Fig. 3F)**. Despite lack of LC *Vegfc* mRNA, incubation of LCs in culture showed release of VEGF-C protein into the supernatant, with less being released from adult LCs **(Fig. 3G).** This result suggested the possibility that LCs could bind VEGF-C expressed by other cells. Indeed, LCs from both young and adult mice bound recombinant VEGF-C, in contrast to T cells and keratinocytes that showed no binding at all **(Fig. 3H)**. Together, these results showed that LCs express PlGF constitutively and suggested that LCs have the potential to promote VEGF-C-mediated functions by binding VEGF-C expressed by cell-extrinsic sources (such as keratinocytes), potentially to deliver to dermal lymphatics.

We asked whether LCs needed to enter dermal lymphatics to exert their effects on lymphatic expansion. CCR7 is essential for LC migration from the dermis into lymphatic vessels (*25*), and adult CCR7-deficient mice showed normal LEC numbers **(Fig. 3I)**. This result suggested that LCs do not need to enter dermal lymphatic vessels to promote lymphatic expansion.

### LC-mediated lymphatic regulation modulates draining lymph node T cell responses to soluble antigen

We assessed the effects of the compromised dermal lymphatics in Lang-DTA mice on lymph node function. As EBD is carried by albumin (*39*), reduced EBD transport to lymph nodes in Lang-DTA mice **(Fig. 1F)** suggested that lymph node responses to soluble antigens would be compromised. Consistent with this idea, adoptively transferred ovalbumin (OVA)-specific CD4 OT-II and CD8 OT-I T cells showed reduced activation and OT-II cells showed reduced proliferation in response to intradermally-injected OVA when in Langerin-DTA hosts **(Fig. 4A-C)**. To understand the extent to which the reduced T cell responses reflected an acute lack of LCs and associated functions such as antigen presentation versus long-term LC effects including LEC development, we depleted LCs in adult Lang-DTR-EGFP mice prior to adoptive transfer of OT-I and OT-II cells. In the absence of LCs but with intact lymphatics **(Fig. 2E)**, OT-II and OT-I T cells did not show reduced responses **(Fig. 4D-F)**. These data suggest that dysfunction of the postnatal LC-lymphatic axis compromises responses to soluble antigen in adulthood.

**Fig. 4.**
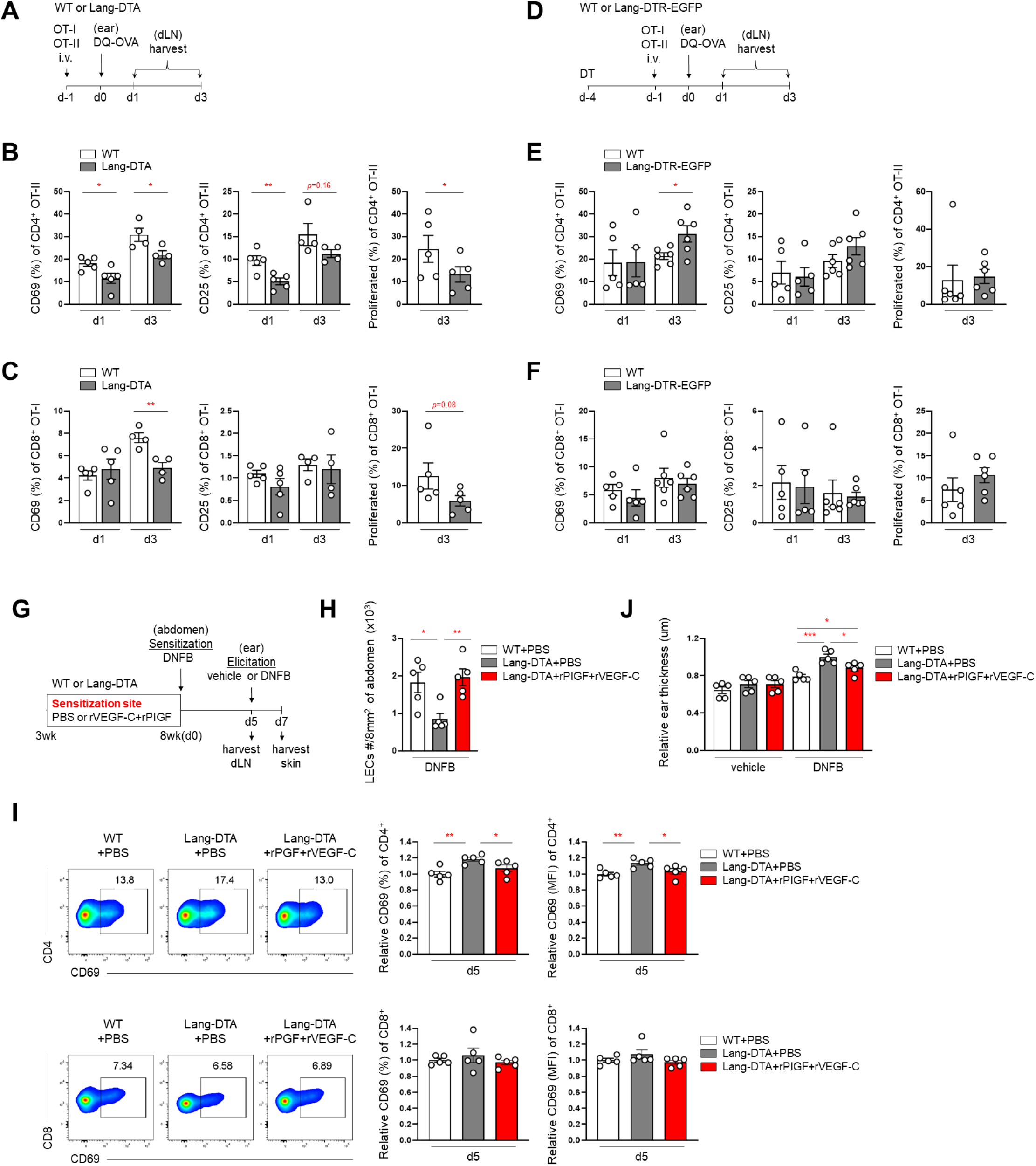
The developmental LC-lymphatic axis regulates T cell responses in adulthood. (**A-F**) CTV-labeled OT-I and OT-II splenocytes were adoptively transferred into WT and Lang-DTA (**A-C**) or DT-treated WT and Lang-DTR-EGFP (**D-F**) hosts one day prior to intradermal OVA on day 0. OTI and OTII T cell responses in draining lymph nodes were examined on days 1 and 3. (**A, D**) Experimental scheme. (**B-C, E-F**) CD69 (left), CD25 (middle), and day 3 proliferation rate (right) of CD4 OT-II T cells (**B, E**) and CD8 OT-I T cells (**C, F**). (**G-J**) 3 week old WT and Lang-DTA mice were treated with PBS or recombinant PIGF+ VEGF-C on shaved abdomen skin until 8 weeks of age. The same skin site was sensitized with 0.5% DNFB on day 0. On day 5, the left ear received vehicle and the right ear received 0.2% DNFB. Inguinal lymph nodes were examined at day 5; and abdominal skin and ear thickness examined on day 7. (**G**) Experimental scheme. (**H**) Sensitization site LEC numbers. (**I**) Lymph node T cell CD69 expression. Representative plots (left), relative percentage (middle) and MFI (right) for CD4 (top row) and CD8 (bottom row) cells. (**J**) Relative ear thickness. Each symbol represents one mouse. Bars represent mean and error bars s.e.m. *p<0.05, **p<0.01, ***p<0.001 using two-tailed unpaired or, for proliferation rate of (**B-C, E-F**), paired Student’s t-test. Mice per condition: n= 4-5 (**B-C**), 5-6 (**E-F**), 4 (**H**), 5 (**I-J**). Data are from 2 (**B-C, E-F, I-J**) and 3 (**H**) independent experiments.

### LC regulation of CHS responses is attributable at least in part to LC-lymphatic axis

We asked if increased CHS responses in Lang-DTA mice (*14*) could be at least in part due to the reduction of LECs. We treated Lang-DTA mice with VEGF-C and PIGF from 3 to 8 weeks of age to restore LEC expansion in the abdominal skin prior to sensitization at this site **(Fig. 4G,H)**. Lang-DTA mice showed increased CD4 but not CD8 activation in inguinal lymph nodes that drained the sensitization site, and LEC restoration with VEGF-C and PlGF reduced the CD4 activation to WT levels **(Fig. 4I)**. The reduced T cell activation was associated with reduced elicitation of CHS responses in the ear **(Fig. 4J)**. These results suggested that LCs regulate CHS responses in part by mediating postnatal dermal lymphatic expansion to modulate T cell priming and that increased CHS responses in Lang-DTA mice reflected in part the loss of lymphatic effects on T cell priming.

## DISCUSSION

Here, we have established an LC-lymphatic axis whereby LCs mediate physiologic postnatal lymphatic expansion and phenotypic changes in the skin. This developmental axis, by facilitating the communication between skin and draining lymph nodes, impacts lymph node T cell responses which, in turn, impacts the control of systemic inflammatory responses as exemplified by CHS responses **(fig. S5)**. This axis is recapitulated during inflammation in adulthood, when LCs function to preserve LEC numbers, suggesting that this axis may serve to promote immune resilience upon skin injury. The results both establish a local skin mechanism for regulating dermal lymphatic function that impacts draining lymph node responses across time and space and point to a mechanism by which LCs control lymph node T cell responses remotely by regulating (skin) tissue development and function.

Our data suggest that LCs mediate postnatal lymphatic development at least in part by expressing PlGF and carrying VEGF-C. LC expression of *Pgf* mRNA was not altered with age, but more VEGF-C could be released from LCs from younger mice compared with adult mice, suggesting the possibility that the specificity of the LC-LEC axis for the postnatal period was in part determined by the level of VEGF-C carried by LCs. LCs had similar capacity to bind VEGF-C at both 4 and 8 weeks, but keratinocytes from 4 week old mice expressed more *Vegfc* mRNA, suggesting that the greater level of LC-associated VEGF-C at 4 weeks of age was due at least in part to greater availability of VEGF-C. VEGFs are well-known to bind heparan sulfates, which regulates their bioavailability and activity (***35, 40–44***); whether LCs bind VEGF-C via heparan sulfate moieties on cell surface glycoproteins remains to be investigated. Together, the data suggest a model whereby, during the postnatal period, LCs in the epidermis acquire VEGF- C from keratinocytes and then carry this VEGF-C to the dermis, where the VEGF-C and LC-derived PlGF can mediate lymphatic expansion by promoting LEC survival **(fig. S5)**. In adulthood, however, keratinocytes express less VEGF-C, reducing LC capacity to regulate LECs until an injury occurs, during which time VEGF-C by keratinocytes or other cell types may be upregulated. The idea of LCs as a ferry for molecules expressed by other cells suggests that LCs serve in part to integrate information about the skin state when regulating lymphatic vessels, with consequent effects on immunity.

Normal dermal LEC numbers in CCR7-deficient mice suggested that LCs can regulate lymphatic expansion without having to enter the lymphatic vessels. Perivascular mesenchymal cells can regulate blood vessel function via VEGF-A (*45*), supporting the idea that LCs can similarly deliver VEGF-C and/or PGF to LECs from the dermis without entering lymphatic vessels. This finding also points to the tissue-modulating functions of LCs that are distinct from antigen-presenting functions, and further studies will be needed to understand to extent to which these different functions reflect the same cells in different states or different subsets of LCs.

Indeed, Ruedl and colleagues recently showed that the dermis contained F4/80^hi^ LCs and another population of cells bearing LC markers CD207, EpCAM, and CD11b that were lower for F4/80 (*46*). The study proposed, based in part on slow partial replenishment by bone marrow-derived precursors, that the F4/80 lower cells were distinct from LCs and, based on lower F4/80 and similar replenishment rates, were the cells that migrated to draining lymph nodes. Although this study did not show definitively that epidermal LCs do not downregulate F4/80 levels and migrate to lymph nodes, it did point to heterogeneity of LC phenotypes in the dermis and the possibility that some LC subsets migrate while others do not. It is possible that the lack of requirement for CCR7 for lymphatic regulation reflects the role of LCs that do not migrate to lymph nodes.

While our efforts here were focused on LC-mediated regulation of lymphatic function, how dermal lymphatic restoration in Lang-DTA mice limits T cell priming during CHS responses remains to be better understood. One possibility is that the dearth of dermal lymphatics in Lang- DTA mice results in reduced skin clearance and greater dermal accumulation of inflammatory mediators, which leads to increased DC maturation and licensing and, consequently, greater lymph node T cell priming. In addition to tissue clearance functions, LECs can directly promote the function of tolerogenic immune cells (*47–50*), and the possibility that greater T cell priming reflects a dearth of tolerogenic cells remains to be examined.

Our findings have potential implications for better understanding disease. The ability of the LC-lymphatic axis to serve as distal regulators of lymph node T cell function across time and space may lead us to reconsider the extent to which adult immune and inflammatory conditions may in part reflect experiences that impacted the LC regulation of lymphatic development in childhood. Partial loss of LCs from the skin occurs with UVR exposure (*51*) , inflammatory cytokines such as TNF and IL1b (*52*), and mechanical stress (*53*), suggesting the possibility that conditions such as chronic sunburns, chronic excoriation, or loss of skin from burns in early life could potentially lead to immune dysregulation in adulthood. Similarly, lupus is characterized in part by reduced LCs in the skin (*18, 54, 55*), raising the possibility that childhood lupus continues into adulthood in part because a disrupted LC-lymphatic axis during childhood leads to chronic immune dysregulation. Similarly, loss of LCs during adulthood and with aging (*56*) could compromise recapitulation of the LC-lymphatic axis during inflammatory injury in adulthood, potentially also leading to immune dysregulation. These possibilities suggest that restoring the LC-lymphatic axis could be a means to treat autoimmune and inflammatory disease.

## ACKNOWLEDGMENTS

The authors gratefully acknowledge the members of the Lu, Santambrogio, and Outzz-Reed labs for helpful discussions, the Weill Cornell Flow Cytometry Core for cell sorting, Anna Benjamin and Michael Glickman for OT-I and OT-II mice, and Melody Zeng for placental tissue.

## Author contributions

Conceptualization and data curation: TTL, JHS Formal analysis: JHS, TTL, RB

Funding acquisition: TTL

Investigation: JHS, RB, ZHF, SC, WDS, RK Methodology: JHS, RB, ZHF, SC, WD Project administration: TTL

Supervision: TTL, LBI, BJM

Writing – original draft: JHS and TTL Writing – review & editing: All authors

## Data and Materials availability

All data are available in the manuscript or the supplementary materials.

## LIST OF SUPPLEMENTARY MATERIALS

Materials and Methods Figures S1-S5

## SUPPLEMENTARY MATERIALS

Materials and Methods Figures S1-S5

## MATERIALS and METHODS

### Mice

Mice between 1-8 weeks old were used unless otherwise stated and were sex and aged matched. Both male and female mice were used for experiments. C57BL/6J, Langerin-DTA, Langerin-DTR, and CCR7-deficient mice, OT-I and OT-II mice were originally from Jackson Laboratory (JAX) and bred at our facility. The WT mice used in experiments involving Langerin-DTA mice were littermate controls obtained by breeding hemizygous Langerin-DTA males and WT females. Langerin-DTR mice were obtained by breeding homozygous pairs. All animal procedures were performed in accordance with the regulations of the Institutional Animal Use and Care Committee at Weill Cornell Medicine (New York, NY).

### Mouse treatments

For in vivo blocking experiments, PBS or Isotype control (5 ug/each ear, Invitrogen, #16-4301-85), anti-PIGF-2 (5 μg/each ear, R&D systems, #MAB465-500), anti-VEGFR1 (5 μg/each ear, R&D sysems, #MAB4711), anti-VEGFR2 (5 μg/each ear, Bioxcell, #BE0060), and anti-VEGFR3 (5 μg/each ear, BioLegend, #140902) were injected daily into bilateral WT mice ears in various combinations using a Hamilton syringe (Hamilton, 10 ul gastight syringe #7653-01, 30-gauge needle #7803-07) from day 21 (3 weeks old) to day 35-42 (5∼6 weeks old) for every day. For in vivo rescue experiments, PBS control, recombinant PIGF-2 (0.5 μg/each ear, R&D systems, # 465-PL-050), and recombinant VEGF-C (0.5 μg/each ear, BioLegend, #775106) were injected into Lang-DTA mice bilateral ear every 2 days from day 21 (3 weeks old) to day 35 (5 weeks old). For CHS experiments, the same recombinant proteins were injected intradermally to shaved abdomen skin of Lang-DTA mice from day 21 (3 weeks) to day 56 (8 weeks) of age.

For diphtheria toxin (DT) treatments, WT or Lang-DTR mice were given DT (250 ng DT/dose, Enzo Life Sciences) intraperitoneally once a week from day 21 (3 weeks) to 8 weeks of age, or from 8 weeks of age to 16 weeks of age.

For UVR treatments, mice were exposed to 4000 J/m^2^ UVR using a bank of four FS40T12 sunlamps, as previously described ^18^ and examined 7 days later.

### Flow cytometry, cell sorting, and quantification

For staining of murine whole skin, single cell suspensions of skin were generated as previously described ^18^. Ear or 8mm^2^ biopsy punch shaved abdomen skin was excised, finely minced, digested in collagenase type II (616 U/mL; Worthington Biochemical Corporation), dispase (2.42 U/mL; Life Technologies), and DNAse1 (80 μg/mL; Sigma-Aldrich), incubated at 37°C while shaking at 100 rpm for 30 minutes, triturated with glass pipettes, and filtered.

For staining of lymph node stromal cells, single cell suspensions were generated as previously described ^26^. Lymph nodes were harvested, minced, and digested with type II collagenase (616 U/mL) and DNAase1 (40 μg/mL), incubated at 37°C for 30 minutes while shaking at 50 rpm, triturated with glass pipettes, and filtered.

Cells were counted using a Multisizer 4e Coulter Counter (Beckman Coulter). To calculate absolute cell numbers per tissue, the percentage of the total of a particular population from flow cytometry analysis was multiplied by the cells per tissue count from the Coulter Counter.

For intracellular staining, BD Cytofix/Cytoperm kit was used for CCL21, eBioscience™ Foxp3/Transcription Factor Staining kit was used for Ki-67.

For sorting of LCs, T cells, and keratinocytes, ear was excised and trunk skin was depilated by shaving and Nair treatment, excised, and subcutaneous fat scraped off. Tissue was incubated with dispase (2.42 U/mL) at 37°C for 60 minutes. The epidermis was then gently peeled, finely minced, digested in type II collagenase (616 U/mL) and DNAseI (80 μg/mL), incubated at 37°C while shaking at 100 rpm for 30 minutes, triturated with glass pipettes, and filtered. Stained DAPI and gated for LCs (CD45+EpCAM+CD11c+), T cells (CD45+CD3+), keratinocytes (CD45-) then sorted using a BD Influx.

For crawl-out cell isolation from epidermal sheets, ear was incubated with dispase (2.42 U/mL) at 37°C for 60 minutes and epidermal sheets gently peeled from the dermis. The sheet was floated in culture (RPMI with 10 % FBS) in 24 well plate for 2 days, and then cells in the media were collected for VEGF-C binding assay.

For T cells analysis, skin-draining auricular or inguinal lymph nodes were collected and mashed through a 70um strainer, and stained with CD3, CD4, CD8, CD69 and CD25.

For flow cytometry analysis, cells were analyzed using FACS Symphony A3 (BD Biosciences) and FlowJo Software (Tree Star).

### LC-LEC co-cultures

C57BL/6 mouse primary dermal LECs (Fisher Scientific, #NC1631656) were purchased and cultured using endothelial cell medium kit (Cell Biologics, #50-104-8393). LECs were plated in 48-well plates (BD falcon, #353078) at 3 x 10^5^ cells per well x 1 day before media was changed to endothelial cells medium without growth factors. At the same time, LCs were added at 3 x 10^4^ per well along with anti-PIGF (10 μg/ml, R&D systems, #MAB465-500), anti-VEGFR3 (5 μg/ml, BioLegend, #140902), or combination. Cultures were harvested 3 days later using Trypsin-EDTA (Corning, #25-053-CI) and CD45-CD31+PDPN+ LECs were assessed.

### LC culture for supernatant collection

Sorted LCs were cultured with 3 x 10^4^ cells per well in 96-well plate (BD falcon, #353072) for 2 days. Culture supernatants were analyzed for VEGF-C using a VEGF-C ELISA kit (CUSABIO, #CSB-E07361M, Houston, TX).

### VEGF-C binding assay

Cells isolated from crawl-out epidermal cultures were washed with PBS and rested in serum-free media with 2 x 10^5^ cells per well in 96-well assay plate (BD falcon, #353910) for 30 min. The media was changed with new serum free media and incubated with his tagged recombinant mouse VEGF-C (10 μg/mL) for 30 min before staining in PBS with anti-his PE (1:100, BioLegend, #362603) and cell surface markers.

### In vivo OT-I and OT-II T cell responses

Splenocytes from OT-I and OT-II mice were labeled with Cell Trace Violet (Invitrogen, #C34557) and 5 x 10^6^ cells were adoptively transferred by retro-orbital injection in 200 ul of PBS. The next day, 20 ng of DQ^TM^ Ovalbumin (OVA) (Thermo Fisher, #D12053) in 1 ul of PBS was injected intradermally into ear skin using a Hamilton syringe (Hamilton, 10 ul gastight syringe #7653-01, 30-gauge needle #7803-07). DQ^TM^ Ovalbumin will fluoresce green when taken up and proteolytically degraded during dendritic cell antigen processing, but was undetectable at the quantity used. Draining auricular lymph nodes were collected at 1 and 3 days later.

### 2,4-dinitrofluorobenzene (DNFB*)-*induced contact hypersensitivity (CHS) model

Mice were sensitized with 25 μl of 0.5% DNFB (Sigma, #D1529) in acetone:olive oil (4:1) onto shaved abdomen skin at day 0. On day 5, CHS response was elicited with 10 μl of vehicle to each side of the left ear and 0.2% DNFB to each side of the right ear. Ear swelling was measured 48 hr after using a pocket thickness gage caliper (Mitutoyo, #037248).

### Real-time quantitative PCR

RNA was extracted from sorted cells using RNeasy plus mini kit (Qiagen, #74134) and cDNA was generated using iScript cDNA Synthesis kit (Bio-rad, #1708891). Target genes amplification was performed with Maxima SYBR Green/ROX Master Mix (Thermo Scientific, #K0222) on a Applied Biosystems^TM^ QuantStudio 6 Pro (Thermo Fisher Scientific, #A43180). Transcript levels were calculated relative to controls and the relative fold change was calculated using the 2^−ΔΔCt^ algorithm. Mouse primer sequences used were: Pgf: 5′-TGCTGGTCATGAAGCTGTTC-3′ and 5′-GGACACAGGACGGACTGAAT-3′ Vegfc: 5′-GGGAAGAAGTTCCACCATCA -3′ and 5′-ATGTGGCCTTTTCCAATACG-3′ Gapdh: 5′-ATGTGTCCGTCGTGGATCTGA-3′ and 5’ -TTGAAGTCGCAGGAGACAACCT-3′

### Wholemount immunofluorescence staining and analysis

For whole mount immunofluorescence staining, ear pinnae were depilated using Nair and fixed with 4% paraformaldehyde (PFA) for 2 hours at 4’c. Anterior and posterior sides were then separated and cartilage removed from the anterior side. Tissues were permeabilized and blocked with Triton-X (0.2%) and BSA (1%), and stained as indicated.

For lymphatic vessel characterization, whole mount ear skin was stained using antibodies to CD31 (Biolegend, #102502), LYVE-1 (R&D, #FAB2125P), and PDPN (abcam, #ab11936). The whole ear was imaged as a Z-stack (4 stacks, 15µm apart) at 20x magnification using a Axioscan 7 (Zeiss) whole slide imager at the HSS Musculoskeletal Tissue Analysis Core. A semi-automated machine learning segmentation tool in QuPath {Bankhead, 2017 #7968] was utilized to segment both Lyve1+ and Lyve1-PDPN+CD31+ vessels in a fixed region of interest (6mm^2^). Percent vessel+ area of total was calculated and the binary masks of vessel positive area were exported to Amira (Thermo Scientific) for network analysis using the centerline tree tool.

Total nodes, branching nodes and segments were calculated from the generated network. The diameter of six vessels (>200 um in length) per ear were manually measured by placing >8 transverse lines perpendicular to the direction of the vessel.

For activated Caspase-3 expression in skin, ear was stained with CD31 (Biolegend, #102502), LYVE-1 (R&D, #FAB2125P), and active Caspase-3 (R&D, #AF835-SP). Tissue was imaged using a LSM880 with Airyscan and FAST Airyscan High Resolution and 32-channel spectral Detectors (Zeiss; Munich, Germany) in the Optical Microscope Core of Weill Cornell Medicine and analyzed using Zen (3.5 blue edition, Zeiss) and Image J software (NIH).

### Lymphatic flow assays

2% Evans blue dye (Sigma-Aldrich, #E2129) solution was injected in into ear pinna (1ul volume) or foot pad (10ul volume) using a Hamilton syringe (Hamilton, 10 ul gastight syringe, 30-gauge needle). Ears were photographed immediately 0 minute or 22 hours later. Ear-draining auricular lymph nodes and footpad-draining popliteal nodes were harvested 2 and 30 minutes after injection, respectively. For EBD retention analysis, ear was harvested 22 hours after injection. EBD was extracted from tissues by incubating in formamide at 56°C temp for 24 hours and quantitated by measuring absorbance at 620 nm (Infinite 200 PRO spectrophotometer (Tecan, Mannedorf, Switzerland)) and calculating against a titration curve.

### Statistical analyses

For figures showing normalized values, the control sample was set to 1, and the experimental samples were normalized relative to the control for that experiment. For experiments that contained more than one control sample, the mean was obtained for the control samples, and the individual control and experimental samples were calculated relative to this mean. Data are presented as mean ± s.e.m. Statistical analyses were performed using GraphPad Prism v9.2.0 software (GraphPad) using two-tailed unpaired or paired Student’s t-test; p<0.05 was considered significant.

---

**figure S1.**
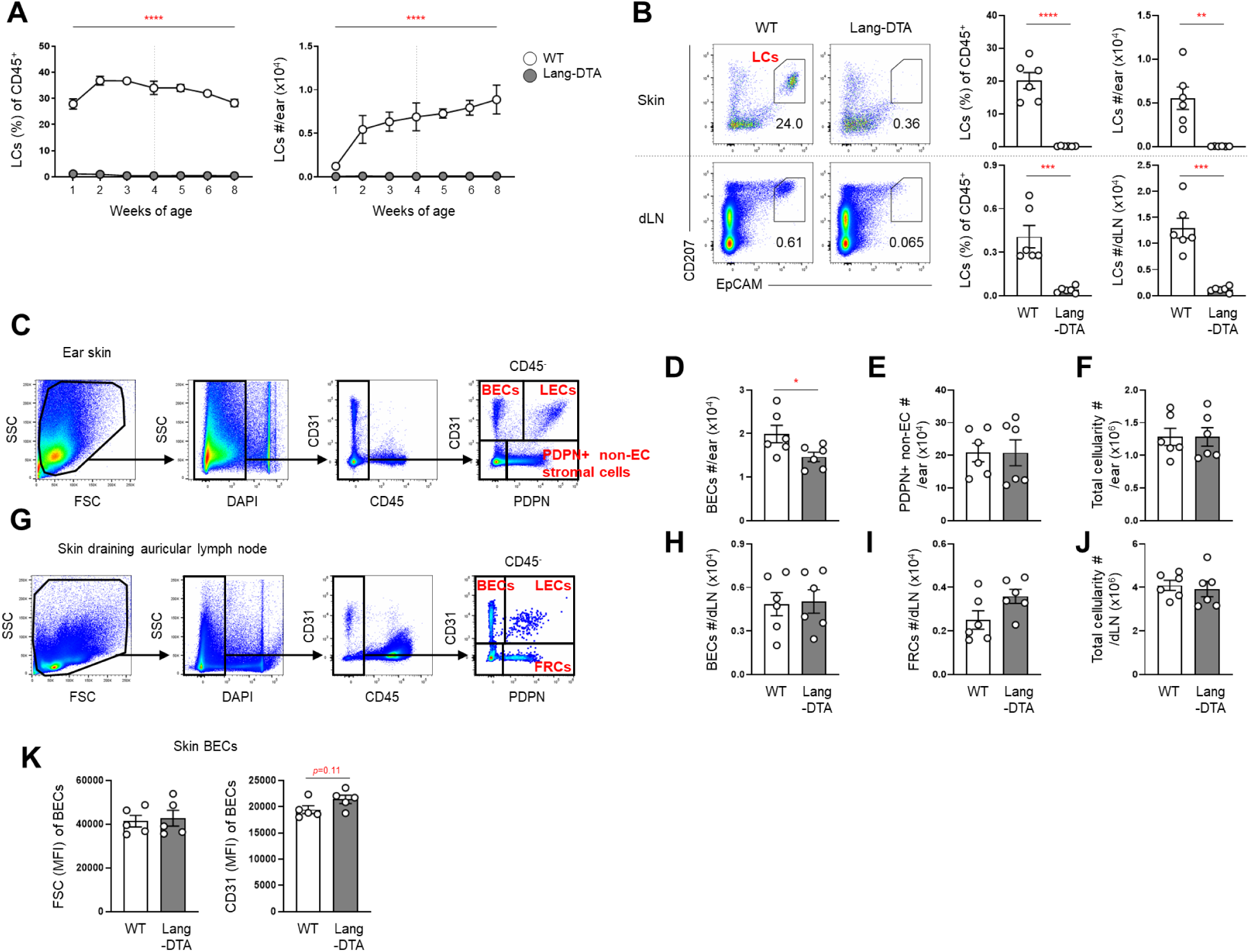
LC and stromal cell numbers in skin and lymph nodes of adult WT and Lang-DTA mice. **(A)** Ear skin CD45+Epcam+CD207+ LC quantitation in WT and Lang-DTA mice at indicated weeks of age, as percentage of CD45+ cell (left) and absolute numbers (right) (*n*=4-7 mice). **(B)** Ear (top) and draining lymph node (bottom) LC quantitation in 8 week old WT and Lang-DTA mice. Representative flow cytometry plots (left), percentage of CD45+ cells and absolute number (right). (*n*=6 mice). **(C-J)** Numbers of indicated cell types in ear skin (**C-F**) and draining lymph nodes (**G-J**) of 8 week old WT and Lang-DTA mice. Flow cytometry gating scheme of CD45-cells (**C,G**), number of BECs (**D,H**), PDPN+ non-EC stromal cells (**E**), fibroblastic reticular cells (FRCs) (**I**), and total cells (**F,J**). **(K)** MFI of FSC (left) and CD31 (right) in 8 week old mice ear skin BECs. Each symbol represents one mouse. Bars represent the mean and error bars s.e.m. Mice per condition: n=4-7 (**A**), 6 (**B-J**), 5 (K). *p<0.05, **p<0.01, ***p<0.001, ****p<0.0001 using two-tailed unpaired Student’s *t*-test. N=4-7 mice per condition and data are from 3 (**A**) and 4 (**B-K**) independent experiments.

**figure S2.**
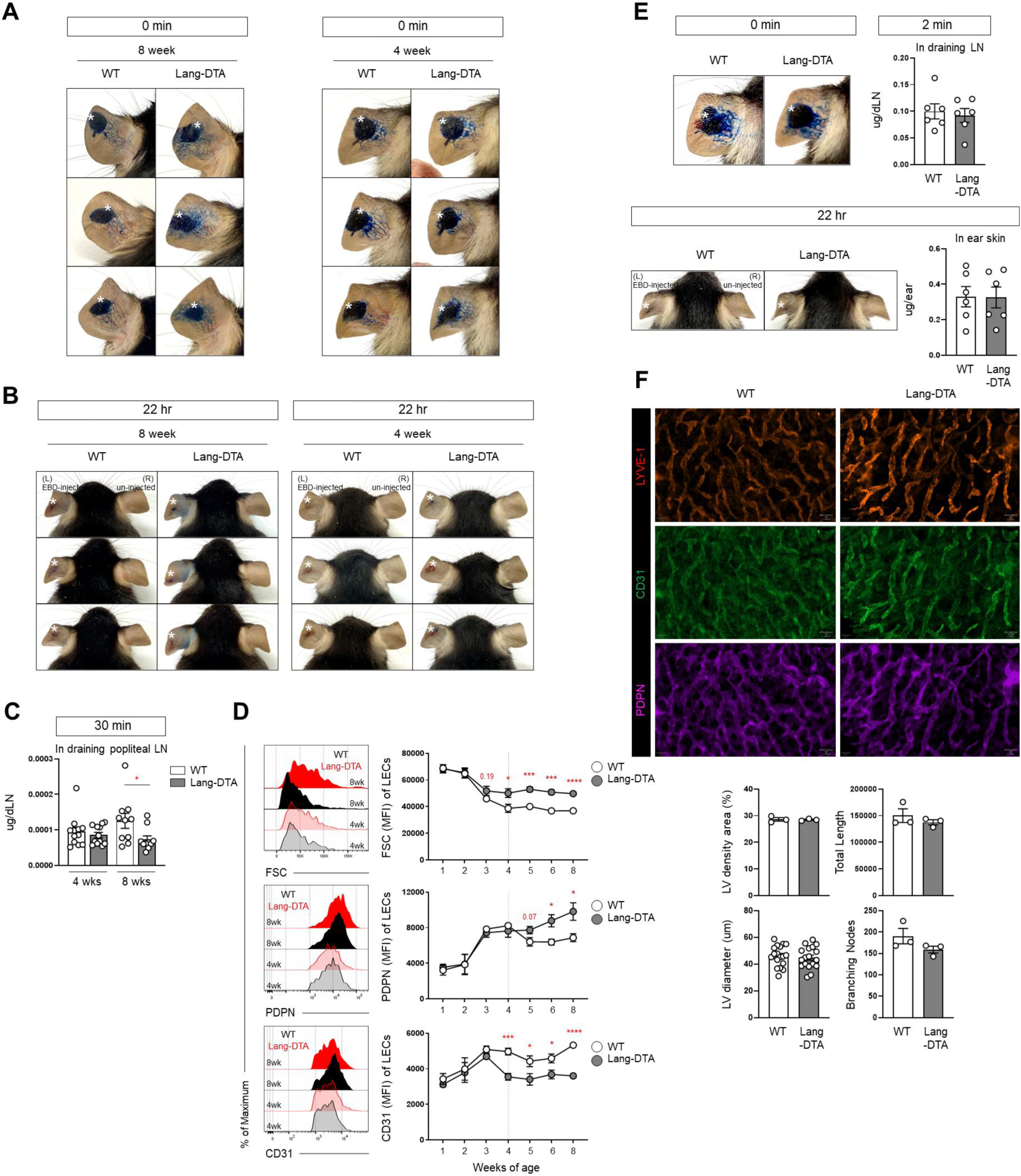
Reduced lymphatic function in Lang-DTA mice at 8 weeks but not at 4 weeks of age. (**A-C**) EBD was injected into ear pinna (**A-B**) or footpad (**C**) of 4 and 8 week old WT and Lang-DTA mice. (**A,B**) Ears were photographed at indicated time points. Stars indicate site of EBD injection. **(C)** EBD quantification in footpad-draining popliteal lymph node at 30 minutes. **(D)** FSC, PDPN, CD31 levels of ear skin LECs at indicated ages. Representative histograms (left), MFI (right). **(E)** EBD was injected into ear pinna of 4 week old WT and Lang-DTA mice and EBD content in draining lymph nodes and ear skin was assessed at indicated times. Stars indicate site of EBD injection. **(F)** Whole-mount staining of 4 week old WT and Lang-DTA ears for LYVE-1 (red), CD31 (green), and PDPN (magenta). Representative images (top), Lymphatic vessel (LV) density, total length, diameter, and branching node numbers (bottom). Symbols represent average of 4-7 mice (**D**); individual segments from images of 3 mice (6 segments per mouse) (**F**, LV diameter); one mouse (**C, F,** LV density, branching nodes, total length). Bars represent the mean and error bars s.e.m. Mice per condition: n=6 (**C,E**), 4-7 (**D**), 3 (**F**). For (**D**), WT and Lang-DTA values were compared statistically at each time point. *p<0.05, **p<0.01, ***p<0.001, ***p<0.0001 using two-tailed unpaired Student’s *t*-test. Data are from 3 (**C-D ,F**) and 4 (**E**) independent experiments.

**figure S3.**
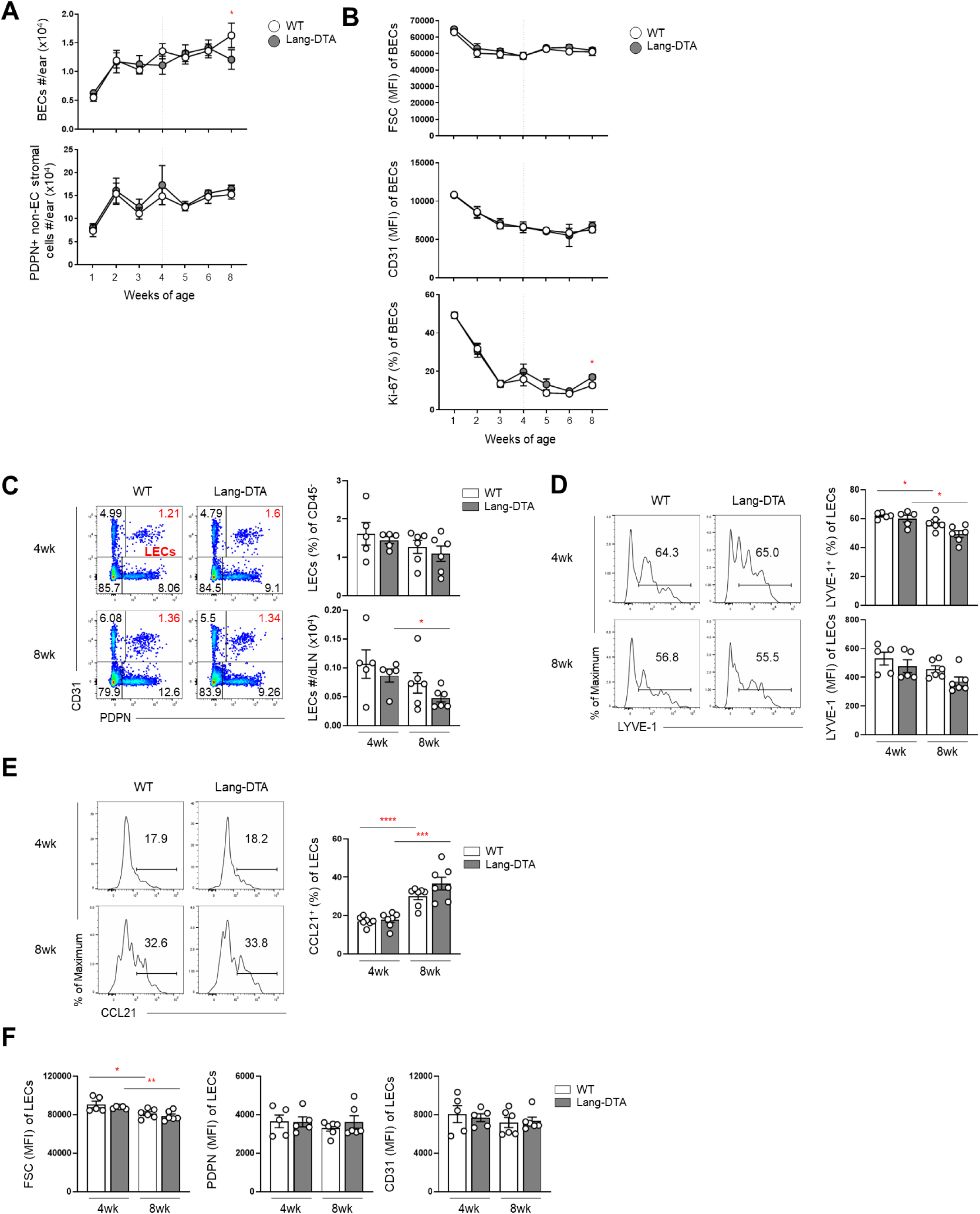
Non-LEC skin stromal cells and lymph node LECs show pattern of postnatal changes that are distinct from that of skin LECs. **(A-B)** Ears of WT and Lang-DTA mice at indicated ages were examined. **(A)** Numbers of ear BEC and PDPN+ non-endothelial stromal cells. **(B)** Ear BEC MFI for FSC (top), CD31 (middle), and Ki-67 (bottom). **(C-F)** Ear-draining auricular lymph node LECs from of 4 and 8 week old WT and Lang-DTA mice were examined. **(C)** Representative plots (left), LEC percentage of CD45-cells and absolute number (right). **(D)** Representative plots (left), percentage of LECs that are LYVE1+ and LYVE1 MFI (right). **(E)** Representative plots (left), percentage of LECs that are CCL21+ (right). **(F)** LEC MFI of FSC (left), PDPN (middle), CD31 (right). Symbols represent average of 4-7 mice (**A-B**) or one mouse (**C-F**). Bars represent the mean and error bars s.e.m. Mice per condition: n=4-7 (**A-B**), 5-6 (**C-F**). *p<0.05, **p<0.01, ***p<0.001,****p<0.0001 using two-tailed unpaired Student’s *t*-test. Data are from 3 (**A-F**) independent experiments.

**figure S4.**
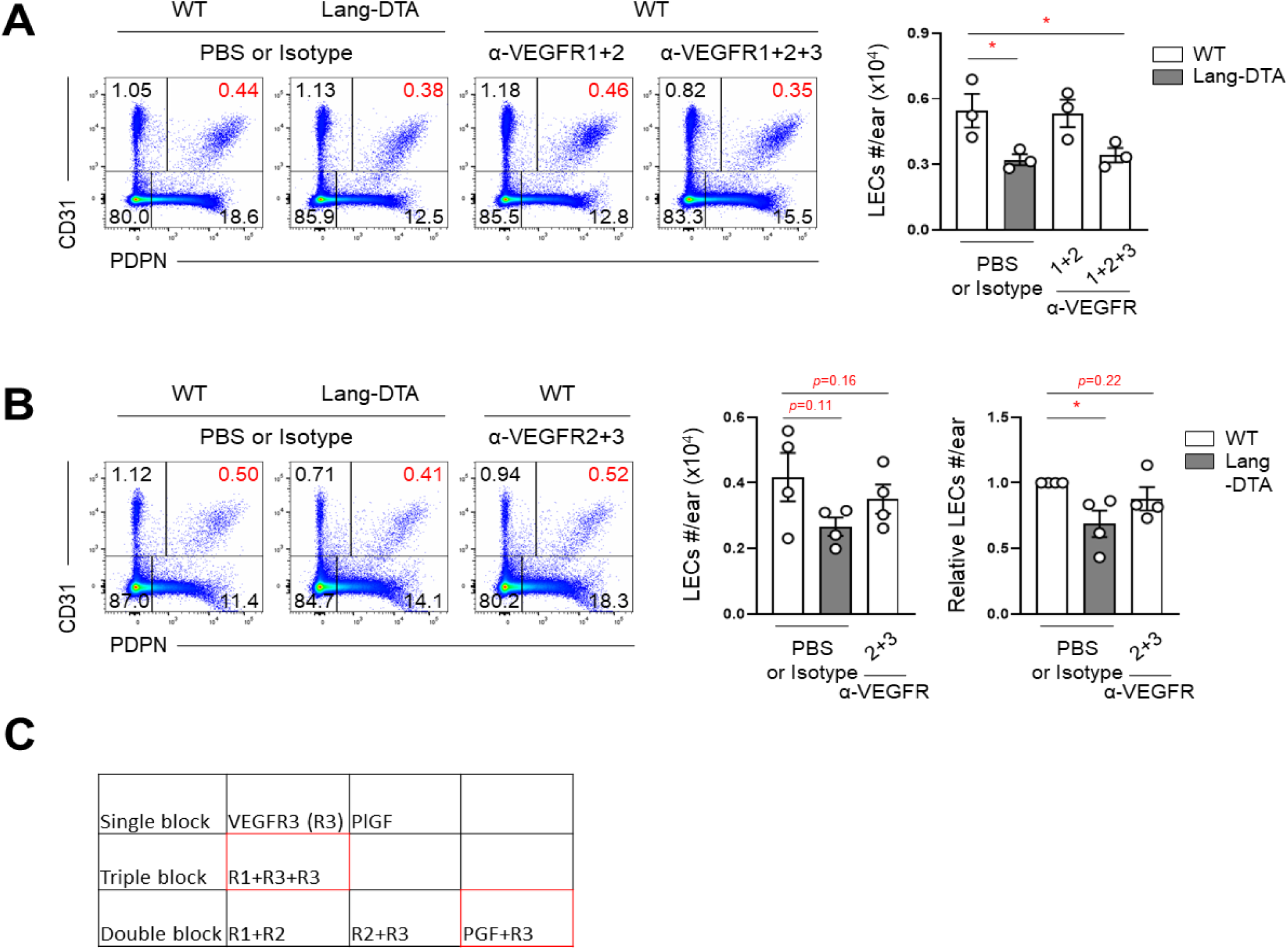
VEGFR1+VEGFR2 and VEGFR2+VEGFR3 blockade have no effect on postnatal lymphatic expansion. **(A-B)** WT mice were treated intradermally with anti-VEGFR1+anti-VEGFR2 or anti-VEGFR1+anti-VEGFR2+anti-VEGFR3 (**A**) and anti-VEGFR2+anti-VEGFR3 (**B**) from 3 weeks of age for 2-3 weeks and then ear LECs quantitated by flow cytometry. Representative plots (left), LEC numbers (right). Each symbol represents 1 mouse. Bars represent mean and error bars s.e.m. *p<0.05 using two-tailed paired (**A, B,** left) or upaired (**B**, right) Student’s *t*-test. Mice per condition: n=3 (**A**), 4 (**B**) and data are from 3 (**A**) and 4 (**B**) independent experiments. **(C)** Summary of antibody blockade experiments shown in **Fig. 3A and fig. S4A-B**. Boxes in red indicate antibody combinations that reduced LEC expansion.

**figure S5.**
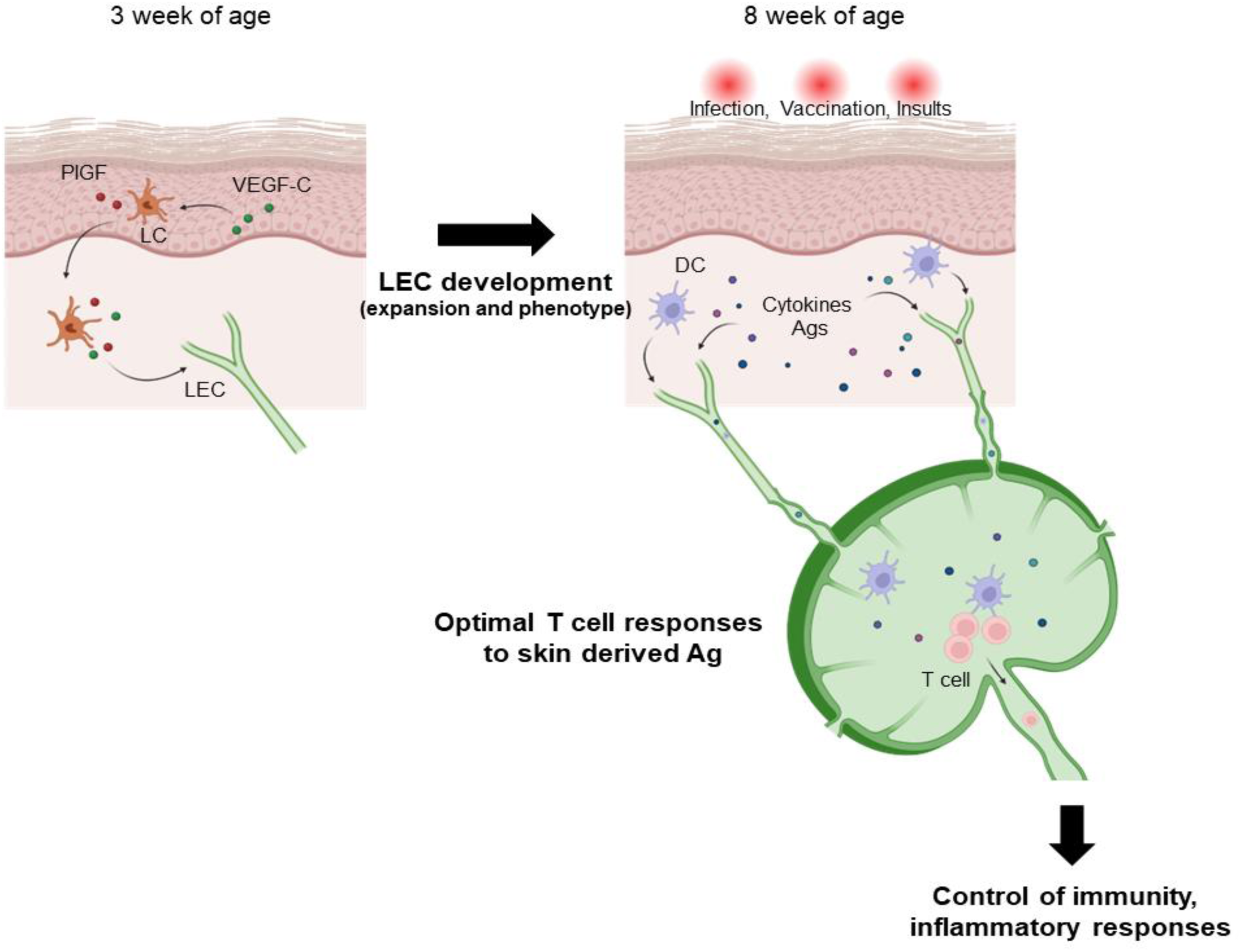
Model of the LEC-lymphatic axis. Between 3 and 8 weeks of age, LCs mediate dermal lymphatic development in part by expressing PGF and carrying VEGF-C. The LC-mediated lymphatic development is essential for normal T cell responses and control of inflammatory responses in adulthood.

